# Phytochemical Analysis and Nematicidal Activity of Medicinal Plants Against *Meloidogyne javanica*

**DOI:** 10.1101/2024.09.29.615697

**Authors:** Saima Maher, Nadra Naheed, Noureen Khan, Hina Ishtiaq, Ghulam Sarwar Solangi, Aasia karim, Muhammad Imran, Erum Iqbal, Farah Mukhtar, Mahganj Bakhshi, Gul Mina, Saleh S Alarfaji

## Abstract

This study examined the nematicidal potential of ethanol extracts derived from four plants, namely *Actinidia deliciosa* (Chev.), *Carica papaya, Citrus paradise*, and *Raphanus sativus*, against the root-knot nematode *Meloidogyne javanica*. Experimental results indicated a significant reduction in the survival of second-stage juveniles of *M. javanica* when exposed to these plant extracts. Notably, concentrations of 2% and 1% were found to be more effective in comparison to 0.5% and 0.25%, yielding statistically significant outcomes. Furthermore, the mortality rate of the nematodes exhibited a direct correlation with the duration of exposure for most plant extracts. The fruit extracts obtained from *A. deliciosa, C. papaya, C. paradise*, and *R. sativus* demonstrated a substantial decrease in nematode infestation compared to the control group. The nematicidal activity of KI3a (BuOH) significant from 0.1356 to 1.9076 over 72 hours, while P-KI-2a (EtOAc) showed rose from 0.1206 to 0.6456. In compare, GF-3 (EtOAc) and Carbofuran to 0.0179 and 0.0098, respectively. PA-1 (MeOH) and RS-3 (EtOAc) showed modest nematicidal potential. This research emphasizes the efficacy of plant extract in mitigating root-knot nematode infestation, offering a promising alternative to chemical managements. Additionally, comprehensive phytochemical analysis should be achieved to recognize the active chemical constituents responsible for their effects. Exploring synergistic effects between these compounds and other natural or synthetic agents could further improve their potential applications in sustainable agriculture, improving crop protection and promoting eco-friendly practices.

## 1. INTRODUCTION

Root-knot nematodes and plant parasitic nematodes, particularly cyst nematodes characterize a major threat to global agriculture, impacting an extensive range of crops comprising fruits, vegetables, and grains. These nematodes are responsible for substantial economic losses they cause approximately 12.3% of crop losses, about $157 billion annually worldwide. (1) The damage caused by these pests not only reduces crop yields but also distresses the quality of agricultural products, leading to further economic impacts for farmers and the agricultural sector. (2) Root-knot nematodes (RKN), particularly those from the Meloidogyne genus, are disreputable for causing major damage to a wide variety of crops, leading to severe economic losses. current chemical treatments for management these pests are increasingly studied due to environmental concerns, prompting the need for alternative, eco-friendly control methods. The organic pesticides have made from botanical extracts are very significance. (3) Numerous plants extracts have revealed that considerable nematicidal activity against the root-knot nematode *M. incognita*. Plants extract from *Nicotiana tabacum, Syzygium aromaticum, Piper betle*, and *Acorus calamus* demonstrating an EC50 five to ten times lower than synthetic pesticides like chlorpyrifos, carbosulfan, and deltamethrin. (4) these plant extracts have proven to be more effective in controlling nematodes compared to chemical nematicides. Under greenhouse environments, the management of plants extracts has demonstrated effective in decreasing nematode infestations, predominantly root-knot nematodes, and increasing crop yields. The aromatic and medicinal plants extracts, are very potential to control nematodes population as they having compounds such as serpentine, phenols, tannins, flavonoids, and cysteine proteinases, have been specially more effective. These plant extracts have been familiar for their anthelmintic properties against animal, human, and plant parasites. (**5)** these botanical extracts have outperformed or confirmed similar efficacy to conventional nematicides. Particularly, the nematicidal activity of *Bidens pilosa* extracts persisted effective even after storage for 12–18 months. (6)

The current research is aim to investigating the nematicidal competences of plant fruits and vegetables extracts against root-knot nematodes, with a specific importance on their effect on Meloidogyne species. The secondary metabolites, such as phenols and flavonoids in kiwi peel, which have previously been related to antibacterial, antioxidant, and potential anticancer properties, (**7)** *Carica papaya* seeds have exposed antimicrobial properties, mainly against both gram-positive and gram-negative microorganisms. Additionally, they grip potential for ulcer treatment and therapeutic. (8) Phytochemicals are vital for eliminating toxins from the body because they possess antioxidant and anti-inflammatory properties. (9) (Table 1). Grapes (*Vitis vinifera*) comprise various secondary metabolites, some of which include known for its potential health benefits resveratrol is an antioxidant that may have cardio-protective and anti-aging properties. These are responsible for the red, purple, and blue colors in grapes. They are potent antioxidants and have been linked to various health benefits. The reported compounds are catechins and epicatechins, which also have strong antioxidant properties and are associated with cardiovascular health. Quercetin is a common flavonol found in grapes and is known for its antioxidant and anti-inflammatory effects. Tannins are responsible for the astringency in grapes and can have antioxidant properties. They also play a role in the texture and taste of wines. These secondary metabolites contribute to the health benefits of grapes and grape-derived products. In *Vitis vinifera* polyphenols, including anthocyanins, flavanols, flavonols, and resveratrol, offer a wide range of medicinal benefits, such as antioxidant, cardioprotective, anticancer, anti-inflammatory, anti-aging, and antibacterial properties. (10) *Vitis vinifera* fruit peel is particularly rich in bioactive substances, including triterpenoids, phenolic compounds, sugars, organic acids, and flavonoids. (11) The cultivation of radish (*Raphanus sativus*) dates back to ancient Asia, where it was used for its therapeutic properties in considering several complaints. (12,13) *Raphanus sativus* is a nutritious powerhouse, comprising nutritive fiber, proteins, carbohydrates, and vital minerals and vitamins. (14) It also harbors unique bioactive components valuable for human health, chiefly glucosinolates. Furthermore, *Carica papaya* leaves have proved considerable nematicidal potential in controlling root-knot nematodes.

Flavonoids have shown their potential as nerve-toxic compounds, effectively reducing the activity of aphids by damaging their spiracles and nerve endings, ultimately leading to their demise. Kiwi fruit is vulnerable to insect and nematode infestations (15) with root-knot nematodes, including *Meloidogyne incognita, M. javanica, M. arenaria*, and *M. hapla*, being significant threats in Pakistan. (16) These nematodes are notorious as polyphagous parasites, well-adapted to parasitizing various plant species worldwide. (17) Additionally, grapefruit and cedar shrubs have exhibited remarkable insecticidal properties, with the citrus-scented oil derived from these plants effectively destroying mosquitoes, ticks, and a wide range of pests. In pursuit of effective and environmentally friendly strategies for combating Meloidogyne root-knot nematode species, this study explores the nematicidal potential of selected medicinal herbs’ ethanol-based extracts, aiming to contribute to the development of successful organic control methods.

## 2. MATERIAL AND METHOD

### 2.1. Collection and Preparation of Samples

*Actinidia deliciosa, Carica papaya, Citrus paradise*, and *Raphanus sativus*, were collected from Saryab Road, Quetta, Pakistan, in November 2020. The fresh fruit samples were first washed thoroughly under running tap water to remove any surface impurities. Subsequently, they were rinsed with distilled water to ensure cleanliness. The ripe fruit samples were carefully dried in the shade for a period of 10-15 days. This drying process was carried out under controlled laboratory conditions. Once the samples were adequately dried, they were ground into fine powders using a grinder. This step helped in obtaining a suitable form for further extraction.

#### 2.1.1. Extraction of Samples

The powdered samples (approximately 40 grams per 500 mL) were subjected to sequential extraction using various solvents, including n-hexane, chloroform, ethyl acetate, methanol, and water. The extraction process was performed using a Soxhlet apparatus and carried out at temperatures ranging from 50 to 60°C, with extraction times varying from 3 to 8 hours. This sequential extraction aimed to separate polar and non-polar fractions in the samples, following the method described by Dane et al. in 2015. (18)

#### 2.1.2. Fractionation of Extract

The methanol (MeOH) extract (approximately 120 grams) of the selected research material was further fractionated using solvents such as hexane, chloroform, *n-*butanol, and water. The chloroform (CHCl3) fractions were subjected to three different pH levels (pH = 3, 7, and 10). In

#### 2.1.3. Collection of Nematode

Meloidogyne spp (Root-knot) nematodes were collected from infested plants. The infected roots were washed, and nematode eggs were extracted using a standard sieving and centrifugation method.

#### 2.1.4. Experimental Procedure

Tomato seedlings were planted in clean soil in separate different test pots. Each pot was inoculated with a specific number of nematode eggs to induce infection. The seedlings were divided into different groups the test treatment were. control (untreated, no extracts), Positive control (treated with a standard nematicide), and Test groups (treated with different concentrations of the prepared plant extracts).

#### 2.1.5. Nematicidal Activity

The method for evaluating the nematicidal activity of methanolic extract of fruit and vegetables against *Meloidogyne javanica* initiates with the growth of a culture of *Meloidogyne javanica* The culture is inoculated on tomato plants (*Solanum lycopersicum*) and maintained under controlled conditions to reduce environmental directly effect. The stock solutions of the methanolic crude extracts were prepared in 5% aqueous DMSO using distilled water to obtain the desired concentrations. Control condition were established, containing a positive control using Furadan at constant concentrations and a negative control using 5% aqueous DMSO. A total no of 100 freshly hatched second-stage juveniles (J2) of *Meloidogyne javanica* were introduced into each treatment group, with the controls. The prepared stock solutions of the test samples are spread into wells of a 96-well plate ensuring that each treatment is replicated effectively (three times) for statistical reliability. The movement of nematodes is observed using a needle under various time intervals, such as at 6, 12, and 24 hours. After each observation period, the living and dead juveniles is counted under the microscope. After a primary experience period of 72 hours, Larvae that show signs of mortality are transferred into a separate cavity filled with distilled water to check their death. Mortality is checked after an additional 24 hours by checking for movement; juveniles that remain immobile are considered dead. The experimental procedure is repeated at least three times to confirm consistency and reliability of results.

Juvenile mortality is calculated using the formula:

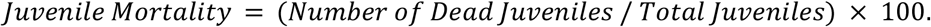

#### 2.1.6. Data Collection

During the experiment, collection of data was accurately collected. These included observations related to plant growth, overall plant condition. The data collected were instrumental in assessing the efficacy of the plant extracts in mitigating nematode-induced damage and promoting the overall health and growth of the tomato plants.

#### 2.1.7. Phytochemical analysis

The phytochemical analysis of the crude methanolic extract derived from selected vegetables and fruits was showed to recognize the potential secondary metabolites present in these plants. Several qualitative tests were performed to identify phytochemicals, **Alkaloids** Detection of alkaloids was carried out using Dragendorff’s reagent. **Flavonoids** Flavonoids were determined using NH3 solution. **Phenolic Compounds** Confirmation of phenolic constituents was performed using Braymer’s reagent. **Carbohydrates** The presence of carbohydrates was tested using Benedict’s reagent. **Tannins** Detection of tannins was carried out with Braymer’s reagent. **Anthraquinones** The presence of anthraquinones was determined using Borntranger’s test. **Saponins** were detected through the formation of foam. **Steroids** were investigated as part of the analysis. **Terpenoids** were assessed using the Salkowski test. These tests provide a comprehensive study about the secondary metabolites present in the methanolic extracts of the selected material, (19)

#### 2.1.8. Biological Assay for *In Vitro* Nematicidal Activity

A biological assay was conducted to evaluate the nematicidal activity of crude extracts obtained from selected fruites and vegetabes extract. The assay involved the following key steps and parameters.

#### 2.1.9. Preparation of Extract Solutions

Five percent methanol-water solutions of the plant extracts were prepared at different concentrations, including 1.0%, 0.5%, 0.25%, and 0.125%.. The experiments were carried out under controlled laboratory conditions. A standard nematicide, furadan, was used as a reference for comparison. Distilled water was employed as the control in the experiments. Each treatment, including the plant extracts and control substances, was replicated twice to ensure the robustness of the results. *Meloidogyne javanica* nematodes were used in the assay. Specifically, 100 larvae were placed in Petri dishes containing 5 ml of distilled water. The Petri dishes were maintained at a constant temperature of 28 ± 2 °C for varying durations: 30 minutes, 1 hour, 2 hours, 3 hours, 4 hours, 24 hours, 48 hours, and 72 hours.

#### 2.1.10 Statistical Analysis of the Experiment

In this research study, the statistical design of the experiment was structured as a completely randomized design. To assess treatment effects, the researchers employed multifactor analysis of variance (ANOVA). When the ANOVA yielded significant results, the distinctions between treatments were further elucidated through Duncan’s multiple range tests at a significance level of P < 0.05. The statistical analysis was facilitated using SPSS statistical software. In the case of percentage data, an arcsine of square root transformation was applied to ensure the appropriateness of the statistical tests. For the analysis of LC50 values, probit analysis was utilized. This analysis was conducted using the PROC PROBIT routine of SAS in the year 2000. By employing these statistical methods and software tools, the researchers were able to rigorously assess the impact of treatments, determine significant differences between treatments, and derive LC50 values, which are crucial in understanding the concentration at which the treatment has a 50% lethal effect. This comprehensive statistical analysis ensured the validity and reliability of the experimental findings.

## 3. RESULT AND DISCUSSION

Furthermore, a comparative analysis was conducted using analysis of variance (ANOVA) to explore the relationships between mortality (%), sample code, and hours (Table 3), as well as between mortality (%), sample code, and concentration (%) (Table 4). Notably, all probability values associated with these analyses were found to be highly significant (P = 0.000) across the various variables examined.

### 3.1.1. One-way ANOVA Analysis

All concentrations examined at various time intervals were subjected to one-way analysis of variance (ANOVA). Table 2 this statistical analysis was performed to assess the differences in mortality rates among the different treatment groups. The results of this analysis are presented in Table 2.

### 3.1.2. Comparison of Means (LSD at P= 0.05)

To further evaluate the differences in mortality rates, the means of mortality rates for each treatment were compared. This comparison was conducted using the Least Significant Difference (LSD) method at a significance level of P= 0.05. LSD is a statistical method that helps identify significant differences between treatment groups.

### 3.1.3. Determination of Median Lethal Concentration (LC50)

The median lethal concentration (LC50) was calculated for the samples administered at various time intervals against root-knot nematodes. LC50 is a crucial metric that indicates the concentration at which a treatment results in a 50% mortality rate among the nematodes. The results of LC50 calculations are presented in Table 3.

#### 3.1.4. General Linear Model Analysis

A general linear model was employed to analyze mortality data in relation to different parameters. This analysis aimed to explore the relationship between mortality rates and various factors or variables. (Table 5).

#### 3.1.5. The statistical software SPSS was utilized to perform this analysis

The results of the general linear model analysis are presented in By subjecting the study’s findings to these rigorous statistical analyses, the researchers ensured a robust evaluation of treatment effects, differences in mortality rates, LC50 values, and the relationship between mortality and various parameters. This approach enhances the reliability and comprehensiveness of the study’s conclusions and insights. (Table 6).

Root-knot nematodes (RKNs) represent a significant and crucial group of plant parasitic nematodes, with *Meloidogyne javanica* being particularly destructive to ornamental and vegetable crops worldwide, notably in regions like Pakistan. *M. javanica* poses a formidable challenge due to its extensive host range, which encompasses over 3000 plant species. ^22, 23, 24^Managing root-knot nematodes is especially challenging because of their subterranean presence, making them difficult for farmers to detect.^25^ The research study revealed significant findings regarding the effectiveness of specific medicinal plant extracts against root-knot nematodes (RKN). The plant extracts were found to possess high levels of toxicity towards RKN, owing to the presence of various secondary metabolites with nematode-inhibiting properties. Among the medicinal plant extracts examined, those of *Withania coagulans, Withania somnifera, Azadirachta indica* (neem), *Tagetes erecta* (marigold), and *Eucalyptus citriodora* (eucalyptus) demonstrated notable nematicidal activity against nematode species including *Meloidogyne incognita, Helicotylenchus multicinctus*, and *Hoplolaimus*. Remarkably, the effectiveness of these plant extracts was comparable to that of chemical nematicide controls. (19)

Furthermore, the research revealed that leaf extracts from *Hunteria umbellate* and *Mallotus oppositifolius* substantially reduced root-knot nematode egg hatching and larval development. This coincided with an enhanced growth of cashew seedlings. (20) In addition, the application of ethanol extracts from *Azadirachta indica* leaves, *Capsicum annuum* fruits, *Zingiber officinale* rhizomes, and *Parkia biglobosa* seeds led to significant improvements in plant height, fruit production, and the weight of *M. incognita-*infected tomato plants compared to non-treated controls. (21) these positive effects were observed consistently across different concentrations. The study also highlighted the rapid nematoxic effect of the plant extracts, which became evident within 24 hours of exposure. While there were significant differences among various concentrations, three specific compounds (PA-L-3a, (EtOAc); KI-2b, (CHCl3); KI-3, BuOH) exhibited remarkable potential, with the ability to kill nematodes at concentrations as low as 0.125%. Similar trends were observed in the LD50 values of these compounds.

Nematodes pose a significant challenge, affecting approximately 15% of all recognized nematode species. They primarily target plant roots, leading to reduced agricultural performance, crop quality, and yields. In Pakistan, the nematode issue is particularly severe and complex due to the country’s tropical climate, which provides a conducive environment for nematode activity and reproduction year-round.

The study’s findings highlight the remarkable efficacy of grapefruit pulp extract in inhibiting the continuous growth of plant pathogens. (Table 1) provides a comprehensive overview of how these extracts exert control over nematode growth. Additionally, kiwi pulp extract demonstrates significant activity, containing secondary metabolites that effectively combat RKNs.

To substantiate these findings, the study subjected the data from various concentrations and time intervals to one-way analysis of variance (ANOVA) (Table 1). The means of mortality rates for each treatment were further compared using the Least Significant Difference (LSD) method at a significance level of P = 0.05. Additionally, the research calculated the median lethal concentration (LC50) for samples administered at different time intervals, targeting root-knot nematodes (Table 1). These analyses collectively underscore the potent effects of medicinal plant extracts from various fruits on RKNs, offering promising strategies for controlling the rapid proliferation of these nematodes. The study sheds light on the substantial potential of plant extracts, particularly from grapefruit pulp and kiwi pulp, in effectively mitigating the growth of root-knot nematodes. These findings hold significant promise for addressing the challenges posed by these destructive pathogens in agriculture and plant cultivation.

## 4. CONCLUSION

Based on the findings of this study, it is evident that fruit extracts obtained from *A. deliciosa* (kiwi), *C. papaya* (papaya), *C. paradisi* (grapefruit), and *R. sativus* (radish) exhibited significant nematicidal potential against *Meloidogyne javanica*, a destructive root-knot nematode. These promising results suggest that these fruit extracts have the capacity to effectively combat this nematode species. However, further research is warranted to delve into the molecular components and bioactive compounds present in these plants responsible for their nematicidal activity. Identifying and isolating these specific compounds will be crucial in understanding the mechanisms behind their effectiveness in controlling M. incognita. Additionally, this knowledge can contribute to the development of more targeted and environmentally friendly strategies for nematode management, particularly in the context of organic farming practices. In summary, while this study provides valuable insights into the nematicidal potential of fruit extracts from certain plants, ongoing research is necessary to unlock the full potential of these natural resources and harness them for effective nematode control in organic agriculture.

## AUTHOR INFORMATION

### Corresponding Authors

**Saima Maher** - Department of Chemistry, Sardar Bahadur Khan Women University, Quetta, Balochistan, Pakistan

**Saleh S Alarfaji -** Department of Chemistry, Faculty of Science, King Khalid University, P.O. Box 9004, Abha 61413, Saudi Arabia

### Authors

**Nadra Naheed**, ICCBS, University of Karachi, Karachi, 72500, Pakistan

**Noureen Khan-** Department of Chemistry, Rawalpindi Women University, Punjab, Pakististan

**Hina Ishtiaq** - Department of Biotechnology, Sardar Bahadur Khan Women University Quetta, Pakistan.

**Ghulam Sarwar Solangi** - Department of Entomology, Shaheed Zulfiqar Ali Bhutto Agricultural College, Dokri, 77060, Pakistan

**Aasia kaim** - Department of Zoology, Sardar Bahadur Khan Women University, Quetta, Baluchistan, Pakistan

**Muhammad Imran**, Department of Chemistry, Faculty of Science, King Khalid University, P.O. Box 9004, Abha 61413, Saudi Arabia

**Erum Iqbal**- National Nematological Research centre, University of Karachi, Pakistan

**Gul Mina** - Department of Chemistry, Sardar Bahadur Khan Women University, Quetta, Balochistan, Pakistan

## Acknowledgement

Authors express appreciation to the Deanship of Scientific Research at King Khalid University Saudi Arabia for funding through research groups program under grant number R.G.P. 2/445/44.

---

**Fig. 1.**
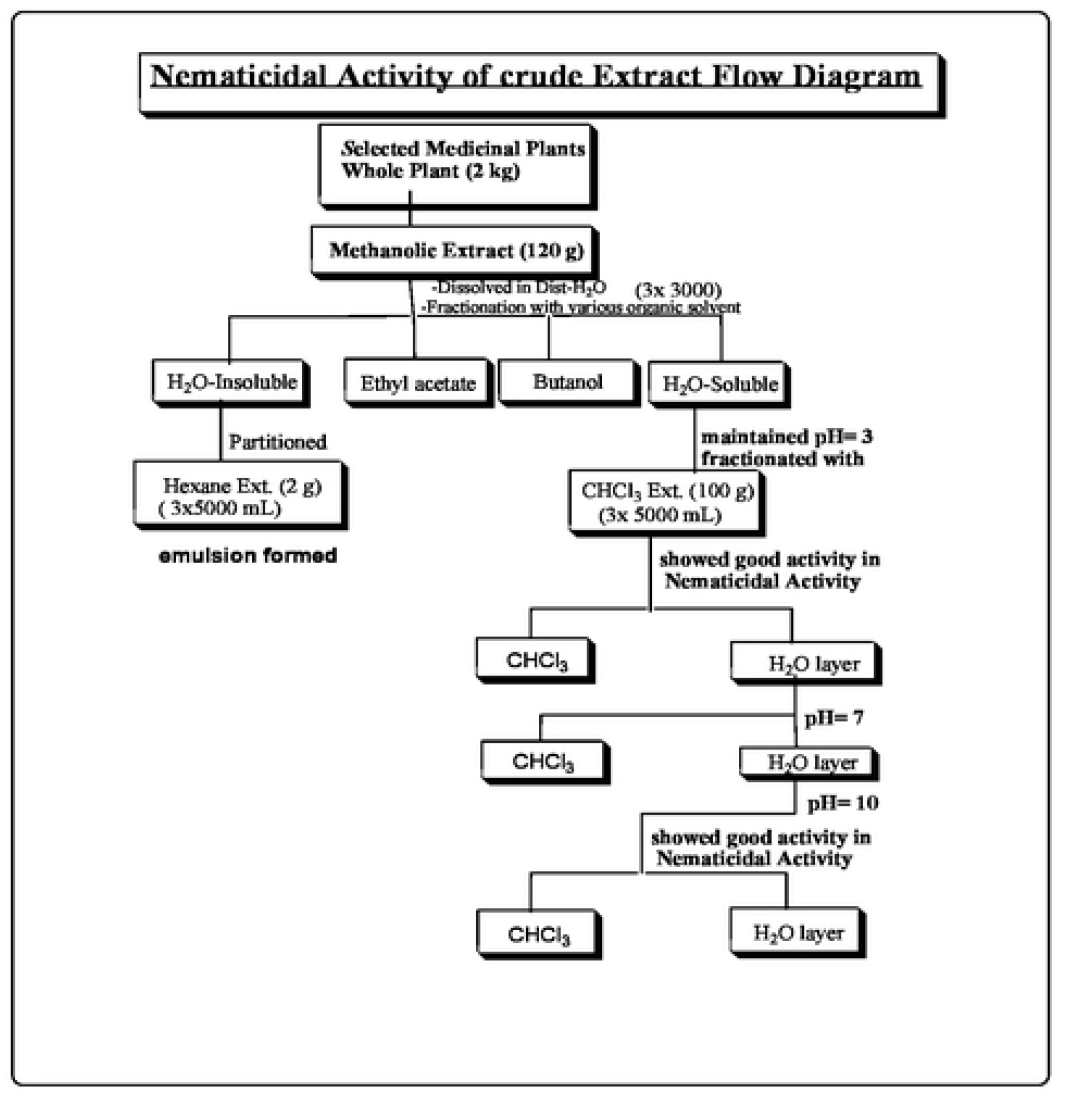
General Fractionation Scheme of Selected Medicinal Plants

**Fig. 3.**
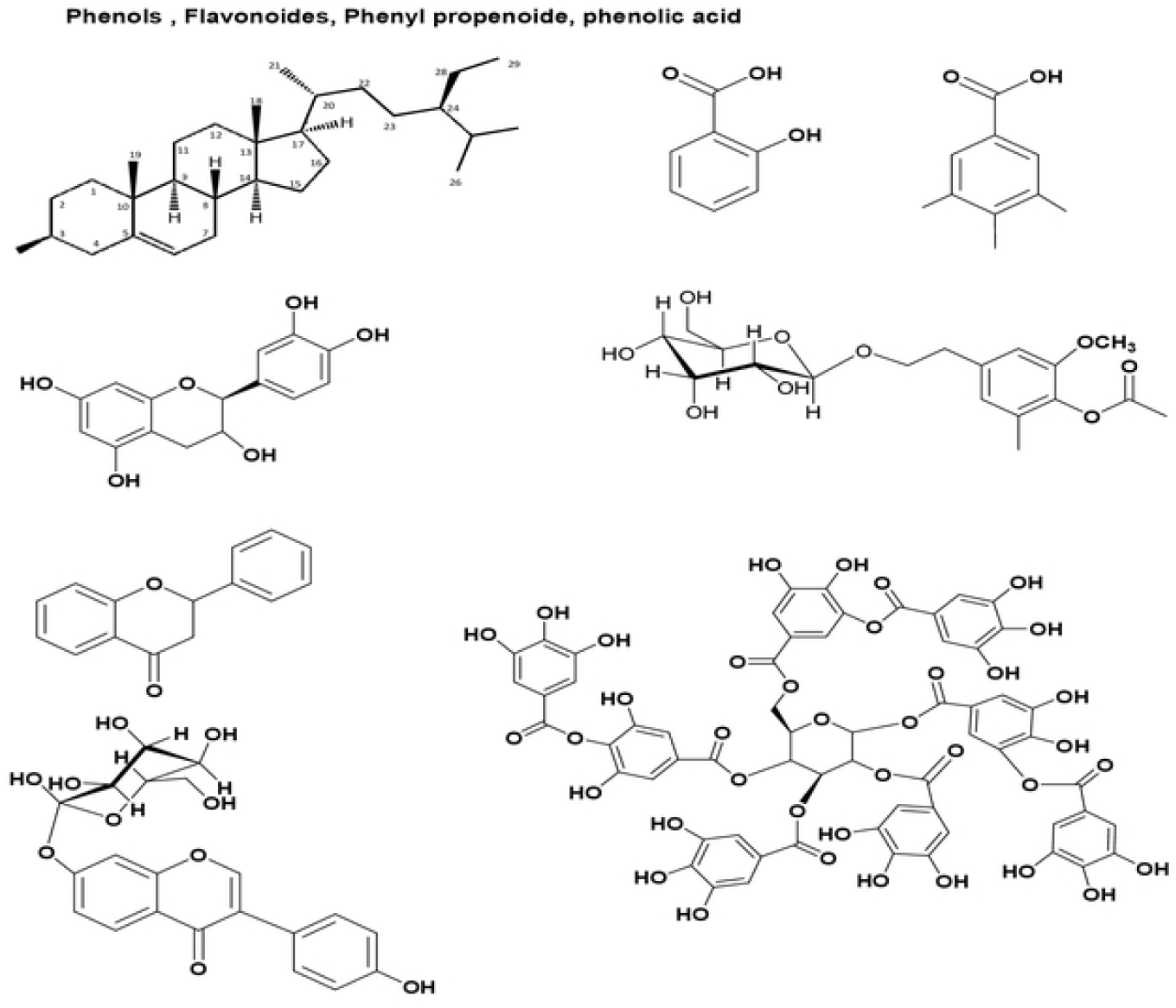
Major secondary metabolites of selected material

**Fig. 4.**
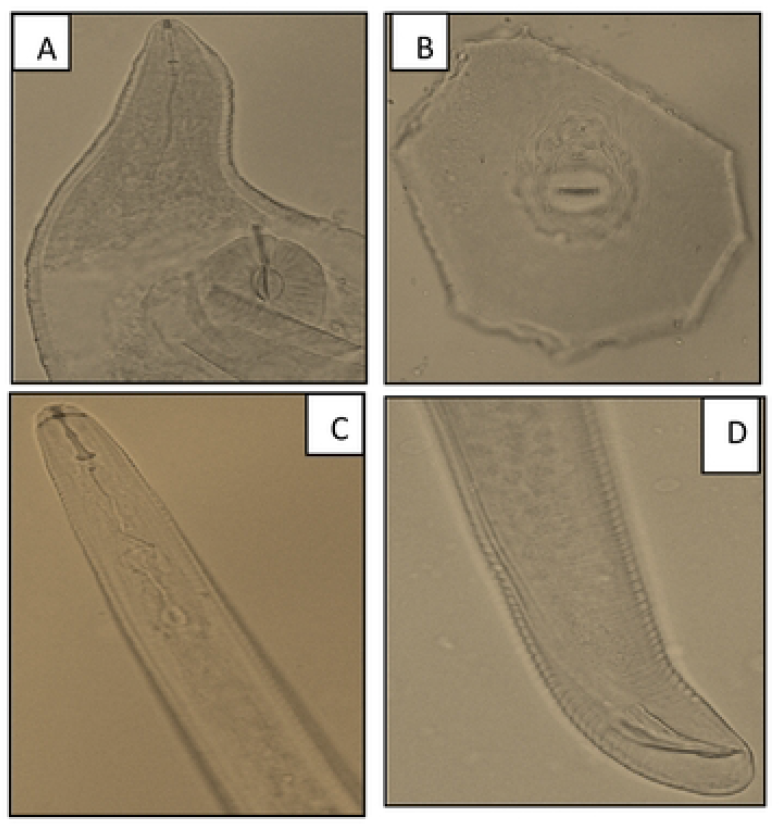
Micrography of *Meloidogyne incognita*.

**Fig. 4.**
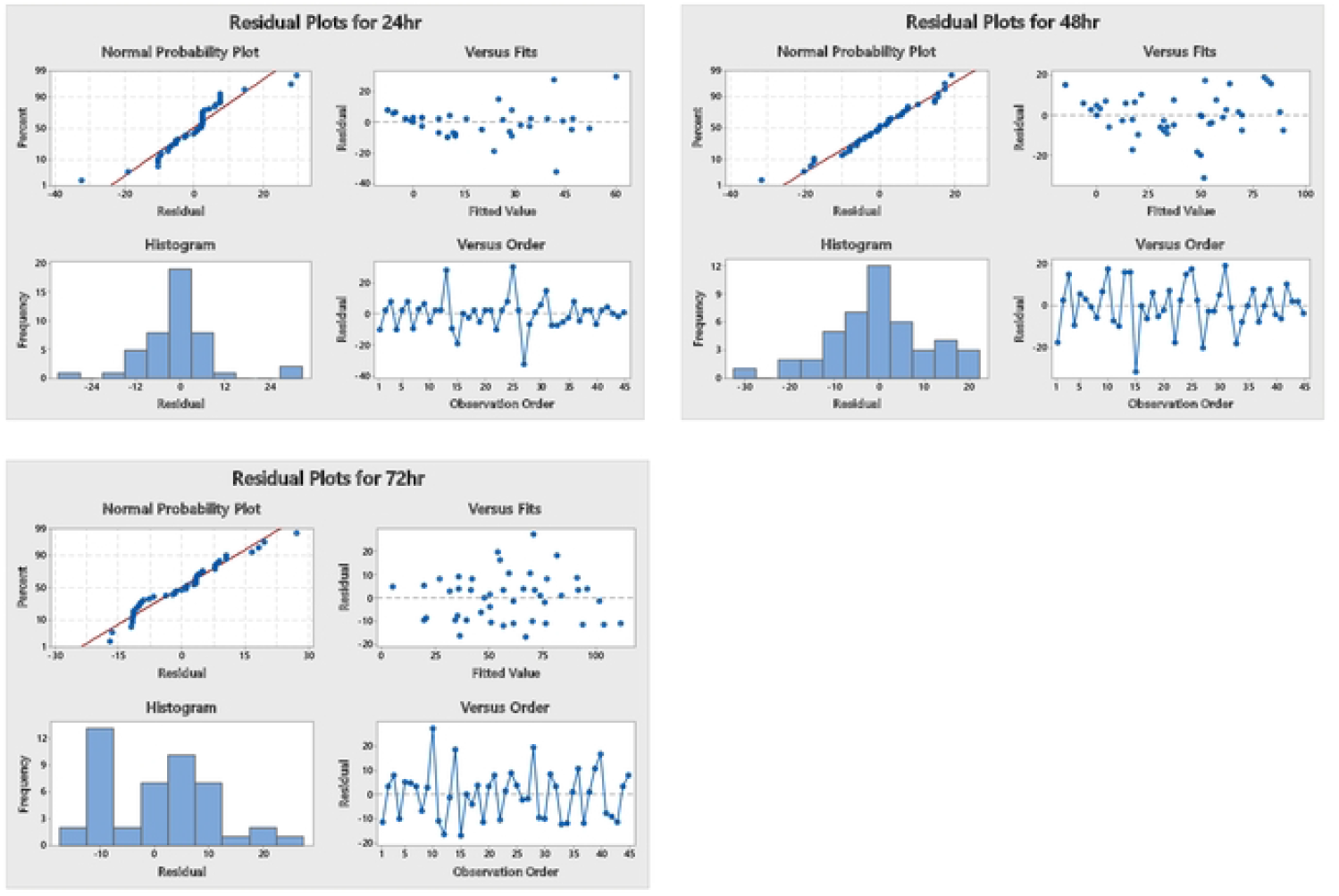
Residual Plots against different time intervals.

